# Beige adipocytes independently improve impaired glucose metabolism in the absence of brown adipose tissues in vivo

**DOI:** 10.1101/2020.06.05.136994

**Authors:** Xiao-wei Jia, Dong-liang Fang, Xin-yi Shi, Tao Lu, Chun Yang, Yan Gao

## Abstract

Beige adipocytes are emerging as an interesting issue in obesity and metabolism research. There is a neglected possibility that brown adipocytes are equally activated when external stimuli induce beige adipocytes. Thus, a question is whether beige adipocytes have the same functions as brown adipocytes when brown adipose tissue (BAT) is lacking. This question has not been well studied. Therefore we determine the beneficial effects of beige adipocytes upon cold challenge or CL316243 treatments in animal models of BAT ablation by surgical denervation. The data show that beige adipocytes partly contribute to impaired glucose metabolism resulting from denervated BAT. Whereas, we found that denervated BAT were activated by cold exposure and CL316243. Thus, we further used BAT-removal animal models to abolish BAT functions completely. We found that beige adipocytes upon cold challenge or CL316243 treatments independently improved impaired glucose metabolism in BAT-removal mice. The insulin signaling was activated in BAT-removal mice upon cold exposure. Whereas, both the activation of insulin signaling and up-regulation of glucose transporter expression were observed in BAT-removal mice with CL316243 treatments. The data show that beige adipocytes induced by cold exposure or CL316243 may have different mechanisms to improve impaired glucose metabolism. Beige adipocytes can also enhance energy expenditure and lipolytic activity of white adipose tissue when BAT is lacking. We provide direct evidences for the beneficial effect of beige adipocytes in glucose metabolism and energy expenditure in the absence of BAT in vivo.

## INTRODUCTION

Obesity is a chronic metabolic disease caused by the long-term energy imbalance. It increases the risk of diseases such as type 2 diabetes, hypertension, hyperglycemia, dyslipidemia and fatty liver (1, 2). The prevalence of obesity presents a significant challenge for public health (1, 2). The current prevention and treatments have not been effective or suitable in the long term (2, 3). Obesity is characterized by increased energy intake and reduced energy expenditure. Therefore, a promising strategy for obesity treatments is to increase energy expenditure by thermogenic adipocytes (4). The thermogenic adipocytes are rich in mitochondria which possess a high level of uncoupling protein 1 (UCP1) expression. When activated, UCP1 dissipates the proton gradient across the mitochondrial inner membrane, uncouples the oxidative phosphorylation and converts the energy of substrate oxidation into heat instead of ATP. This process is referred to as adaptive thermogenesis, a major contributor to total energy expenditure.

In rodents, there are two types of thermogenic adipocytes with high metabolic activity: brown adipocytes and beige/brite adipocytes. Brown adipocytes and beige adipocytes are characterized by multiocular morphology and abundant mitochondrial content containing UCP1 (5, 6). Classical brown adipocytes are mostly distributed in interscapular brown adipose tissue of mice and human infants, and neck brown adipose tissue of human adults (6–13). In rodents, beige adipocytes are recruited and clustered in various white adipose tissues in response to browning stimuli (6, 14). In human adults, beige adipocytes are present in supraclavicular fat of human body (15, 16). The activation of brown adipocytes has been considered an attractive target for therapeutic intervention in obesity and metabolic diseases (17–21). Studies in mouse models have shown that BAT activation can improve hyperlipidemia and harmful effects of obesity (5, 22). Increasing brown fat mass by transplantation also has beneficial effects of decreasing body weight and improving glucose metabolism and insulin sensitivity (5, 23). In addition, several studies have revealed the presence and gene expression of brown adipose tissue in neck and suprascapular areas of human adults. Brown adipose tissue in the human neck area is heterozygous for the coexistence of brown adipocytes and beige adipocytes. Whereas brown adipose tissue from the supraclavicular area are mainly composed of beige adipocytes (15, 16, 24, 25). Aside from BAT, recruitment of beige adipocytes in white adipose tissue (WAT browning) is emerging as an interesting and promising target for therapeutic intervention in obesity and metabolic diseases (3, 5, 6, 17, 21, 26–29). This is due to the fact that beige adipocytes can dissipate energy via UCP1, be recruited and differentiate from mature white adipocytes (30–33) or PDGFRα^+^ progenitors(34), smooth-muscle lineage/mural progenitors (35–37). A recent study revealed that beige adipocytes could differentiate from MyoD^+^ progenitors (38). In human adults, beige adipocytes benefit the insulin sensitivity and the reduction in body weight (39). Additionally, browning in human sWAT in response to cold or Mirabegron (a β3-adrenoceptor agonist) occurs in either obese or older subjects (40).

Cold exposure and β3-adrenoceptor agonist are two traditional and extensively studied physiological stimuli to induce the formation of beige adipocytes in white adipose tissue and activate brown adipose tissue in vivo. Under cold conditions, more norepinephrine (NE) is released by the sympathetic nerve terminals and acts at β3-adrenoceptors. Subsequently, Gas proteins are coupled by β3-adrenoceptors and activate adenylate cyclase (AC) to enhance cyclic AMP (cAMP) levels followed by the protein kinase A activation and phosphorylation of p38 mitogen-activated protein kinases (p-p38MAPK). P-p38MAPK phosphorylates PPARγ cofactor-1α (PGC-1α), cAMP response element-binding protein and activating transcription factor 2 to increase UCP1 protein levels for a greater mitochondrial uncoupling capacity (3, 41–43). Multiple agents such as β3-adrenoceptor agonist and PPARγ agonists are also able to induce browning of white adipose tissue (3, 5, 14, 26, 44)}. In addition to the effect on the formation of beige adipocytes, cold exposure and β-adrenorceptor agonist, as well as PPAR agonists, have been demonstrated to be capable of activating brown adipose tissue in animal models (45–47). This effect is due to the arrangement of the sympathetic nervous system and β-adrenorceptors both in BAT and WAT. Recently, Labbe’s group showed that iWAT did not exhibit significant oxidative activity after adrenergic stimulation in the absence of a functional BAT (48). We asked whether beige adipocytes have the same functions as brown adipocytes, in the absence of brown adipose tissues. The answer to this question has not been thoroughly determined. In the present study, we used surgical denervation and removal of BAT as an ablation system to investigate the functions of beige adipocytes in vivo. We investigated the glucose metabolism and energy expenditure of animal models of BAT ablation after browning stimuli. While we also investigated the glucose transporter genes and insulin signaling in animal models. We aim to determine the beneficial effects of beige adipocytes in the absence of BAT in vivo.

## MATERIALS AND METHODS

### Animals and experimental design

Adult male C57BL/6J mice (6-8 weeks old, 20-25g) were used in this study and housed with a 12-h light/12-h dark cycle. Both water and foods were available ad libitum. The experimental procedure were conducted according to the National Institutes of Health Guide for the Care and Use of Laboratory Animal and approved by the Animal Use and Care Committee of the Capital Medical University, Beijing, China. The study was carried out with the minimum number of animal, and all precautions were taken to avoid animal suffering.

The mice were assigned into seven groups: Group 1 (denoted as DN+CE, n=8): mice received surgical dennervation of BAT and then were kept under cold conditions (4°C) for 14 days. Group 2 (DN+CL, n=8): mice received surgical denervation of BAT and intraperitoneal injections of CL316243 (C5976; Sigma) at 1mg/kg for consecutive 14 days. Group 3 (DN+RT, n=8): mice received surgical denvervation of BAT and were kept under room temperature (around 20°C). Group 4 (RM+CE): mice received surgical removal of BAT and then were kept under cold conditions (4°C) for 14 days. Group 5 (RM+CL, n=8): mice received surgical removal of BAT and then intraperitoneal injections of CL316243 at 1mg/kg for consecutive 14 days. Group 6 (RM+RT, n=8): mice received surgical removal of BAT and were kept under room temperature around 20°C. Group 7 (Sham, n=8): mice received the same surgical procedure without cutting nerve bundles of BAT and removal BAT. Bodyweight and food intake were measured every week. All animals were euthanized three weeks after surgical procedure and prepared for the intraperitoneal glucose tolerance test (IPGTT) assay and intraperitoneal insulin tolerance test (IPITT) before euthanasia. BAT, subcutaneous white adipose tissue (sWAT) and epididymis white adipose tissue (eWAT) were prepared for quantitative Real-time PCR (qRT-PCR), western blot, hematoxylin-eosin (H&E) staining and immunostaining. The BAT-removal mice with CL316243 treatments or not were also monitored for energy metabolism analysis.

### Surgical denervation and removal of BAT

As previous descriptions (49–52), mice received a surgical denervation or removal of BAT. Under a stereomicroscope, a midline incision in the skin was performed along the upper dorsal surface. The BAT pads were separated carefully from the surrounding tissues by blunt dissection. For surgical denervation, the five nerve bundles were cut without damage to the blood vessels to BAT. The denervated BAT and surrounding tissues were kept in the original positions. For surgical removal of BAT, the blood vessels of BAT were dissociated and ligated by absorbable surgical sutures. Then, BAT pads were removed, and the surrounding tissues were maintained to their original locations. The same surgical procedure without cutting the nerve bundles and removal of BAT fats were defined as the sham group. All mice were allowed to recover for one week for the following experiments.

### Energy metabolic Study

For energy metabolism analysis, mice after treatments were monitored using Sable Systems (Sable Systems International, Las Vegas, Nevada, USA). The mice were placed in metabolic cages for 48 hours to acclimate to the environment first, and metabolic data were then collected for 72 hours for analysis. Oxygen consumption (VO^2^), carbon dioxide production (VCO^2^), respiratory exchange ratio (RER), energy expenditure (EE), were measured every 5 min for analysis. Food and water intake were also measured using a precision scale and volumetric drinking monitor, respectively. The ambulatory activity was estimated by the number of infrared beams broken along the x-axis and y-axis of the metabolic cage and the pedometers.

### Glucose tolerance and insulin tolerance test

Glucose tolerance was evaluated using IPGTT in all groups. All mice were fasted for 16 hours with free access to water, and prepared for intraperitoneal injection of glucose (1 g/kg). Blood glucose concentrations were collected at the indicated time points (0, 15, 30, 60, 90 and 120 minutes) after glucose administration and determined by a NovaMax glucometer. Insulin tolerance was measured using IPITT in all groups. Briefly, mice were fasted for 6 hours with free access to water before IPITT assay. Mice were administrated by intraperitoneal injections of insulin at a dose of 0.75 U /kg body weight. Whole Blood was then collected from the distal part of the tail vein at the indicated time points (0, 15, 30, 60 minutes) after insulin administration. Blood glucose concentrations were determined by a NovaMax glucometer.

### Quantitative Real-time PCR

According to the manufacturer’s instructions, total RNA was extracted from the BAT, sWAT and eWAT using the TIANgen RNA extraction kit (Tiangen, China). qRT-PCR was conducted with SuperReal PreMix Plus (SYBR Green) (TIGANGEN, Beijing, China) in a final volume of 20 μl containing 10 μM each of the forward and reverse primers. β-actin was used as an internal control to normalize the results of mRNA in each sample. The sequences of primers were listed in supplementary Table S1. Relative mRNA levels were measured using the CFX96 Real-time System, C1000 Thermal Cycler (BioRad).

### Western blot

The adipose tissues were homogenized, lysed and sonicated in RIPA buffer containing protease inhibitor cocktail on ice. After centrifugation, the supernatant was collected and stored at −80 °C until assay. Protein concentration was determined by BCA protein assay kit (Thermo Scientific, Rockford, IL., USA). Equivalent amounts of protein were resolved by 10% SDS-PAGE and transferred to polyvinylidene difluoride membrane. The membranes were incubated with 5% non-fat dry milk or 5% BSA in TBST and then followed by the primary antibodies and secondary antibodies. The primary antibodies were anti-UCP1 (Abcam, Cambridge, MA, USA), anti-Glut1 (Santa Cruz, Texas, USA), anti-Glut4 (Santa Cruz), anti-TH (Santa Cruz), anti-β-actin (Abcam, MA, USA), anti-p-AKT (Cell Signaling Technology, Beverly, MA, USA), anti-AKT (Cell Signaling Technology), anti-p-IRS1 (Cell Signaling Technology), anti-IRS1 (Cell Signaling Technology). β-actin was used as the loading control. The bands were analyzed by the Image J software (NIH, Bethesda, MD, USA). The protein amounts were normalized to that of β-actin.

### Histology and Immunohistology

BAT, sWAT, and eWAT were collected and then fixed with 4% paraformaldehyde at 4°C for 48 hours. All tissues were dehydrated through serial ethanol concentrations, embedded in paraffin and then sectioned at 6-μm thickness. The sections of BAT, sWAT, and eWAT for each mouse were prepared and stained with H&E and immunohistochemistry. The avidin-biotin complex immunoperoxidase technique was used to visualize immunoreactivity. The primary antibodies were anti-UCP1 (Abcam) and anti-TH (Santa Cruz). Primary antibodies were omitted as a negative control.

### Statistical analyses

All Data are presented as the mean ± standard error of the mean (SEM) and subjected to statistical analysis using GraphPad Prism 6.0 (GraphPad Software, Inc., La Jolla, CA, USA). The data were analyzed by one-factor ANOVA followed by Tukey’s test. Statistical significance was defined at P < 0.05.

## RESULTS

### Denervation of BAT did not alter the body weight and food intake, but impaired glucose metabolism at room temperature

To inhibit the BAT activity, surgical denervation was used currently to remove nerve bundles from BAT. As illustrated by arrows in Figure 1A, the five nerve bundles of each lobe of BAT were exposed and surgically removed. Tyrosine hydroxylase (TH, an initial and rate-limiting enzyme in norepinephrine biosynthesis) was selected for denervation verification of BAT. A number of TH positive fibers were found in BAT in Sham mice. In contrast, a few of TH positive fibers were found in BAT in the denervated mice (Figure 1B). The data for western blot showed that the levels of TH protein were significantly decreased in BAT in Denervation group when compared with that in Sham group (Figure 1C). These results suggest the efficiency of bilateral denervation in BAT.

**Figure 1.**
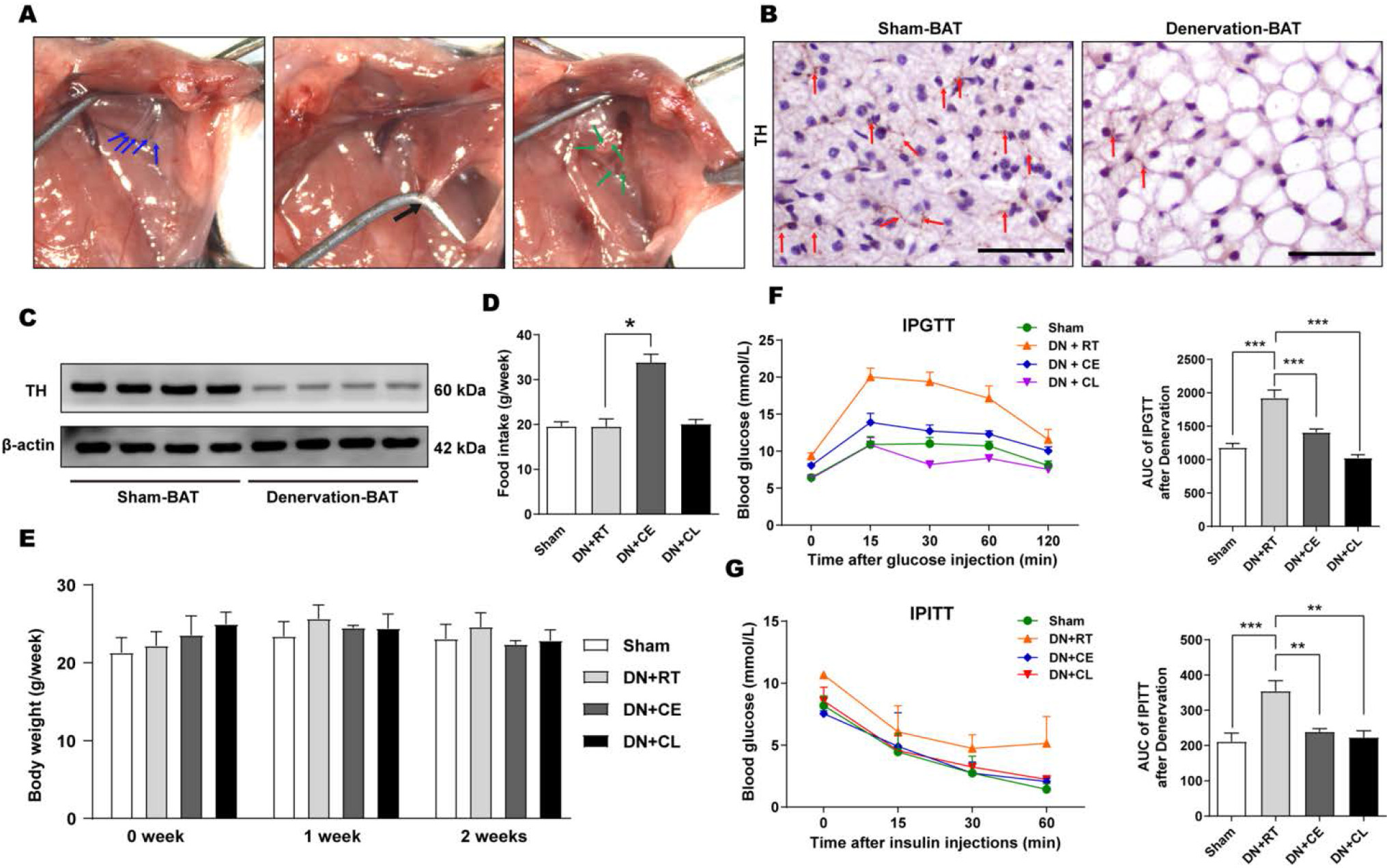
Beige adipocytes partly improved impaired glucose tolerance of denervated mice. (A) Surgical denervation of BAT; The blue arrows showing the intact five nerve bundles (left), the black arrow showing the dissociated nerve bundles (middle) and the green arrows showing cutted nerve bundles of BAT (right). (B) TH immunostaining in BAT in Sham (left) and denervated (right) mice. The red arrows showing TH positive fibers in BAT. Scale bar =50 μm for B. (C) Western blot showing the protein levels of TH in denervated and Sham mice. (D and E) The diagramS showing the food intake (D) and body weight at 0 week, 1 week and 2 weeks (E) of mice in Sham, DN+RT, DN+CE and DN+CL groups. (F and G) The diagrams showing the blood glucose (mmol/L) at indicated time points after glucose injections (F, left), AUC analysis during the measurement (F, right), the blood glucose (mmol/L) at indicated time points after insulin injections (G, left) and AUC analysis during the measurement (G, right) of mice in Sham, DN+RT, DN+CE and DN+CL groups. A one-factor ANOVA followed by a Tukey’s test was used to make group comparisons. *p < 0.05, **p < 0.01, ***p < 0.001 versus indicated groups. BAT, brown adipose tissue; CE, cold exposure; CL, CL316243. AUC, area under curve; DN, denervation; IPGTT, intraperitoneal glucose tolerance test; IPITT, intraperitoneal glucose tolerance test; RT, room temperature; TH, tyrosine hydroxylase.

A one-factor ANOVA indicated that there was no difference in food intake (Figure 1D) and body weight (Figure 1E) between Sham and DN+RT groups. The data for IPGTT showed that the blood glucose levels were significantly higher in DN+RT group (Figure 1F, 9.34 ± 0.46 mmol/L) when compared with that in Sham group (Figure 1F, 6.43 ± 0.35 mmol/L). In addition, the AUC of IPGTT and IPITT were significantly increased in DN+RT group when compared with Sham group (Figure 1F and G). These data suggest that surgical denervation of BAT results in impaired glucose metabolism.

### Beige adipocytes in sWAT and eWAT partly improved impaired glucose metabolism of denervated mice

To determine the ability of beige adipocytes in sWAT and eWAT in denervated mice, cold exposure and intraperitoneal CL316243 (β3-adrenergic receptor agonist) injections were used to induce beige adipocytes. As showed in Figure 2A, sWAT showed a dark brown color in the denervated mice after cold exposure or CL316243 injections. In contrast, eWAT showed a dark brown color in the denervated mice after CL316243 injections, but not cold exposure. The denervated BAT was paler than the innervated BAT. An unexpected result was that denervated BAT also presented a dark brown color in the mice after cold exposure or CL316243 injections. The qRT-PCR analysis showed that the *Ucp1* mRNA level were significantly increased in the BAT of mice in DN+CE and DN+CL groups (Figure 2B). The levels of *Ucp1* mRNA in sWAT were also increased by many folds in DN+CE and DN+CL groups when compared with that in DN+RT and Sham groups (Figure 2B). Whereas the levels of *Ucp1* mRNA in eWAT were significantly increased by many folds in DN+CL, but not in DN+CE groups (Figure 2B). The significant levels of UCP1 protein were detected in BAT in DN+CE and DN+CL groups (Figure 2C). UCP1 protein levels were also increased by seven folds in sWAT in DN+CE and by five folds in DN+CL groups. No UCP1 protein was detected in eWAT in Sham group (Figure 2C). There was a significant difference in UCP1 protein levels in eWAT in DN+CL group when compared with DN+RT and DN+CE groups (Figure 2C).

**Figure 2.**
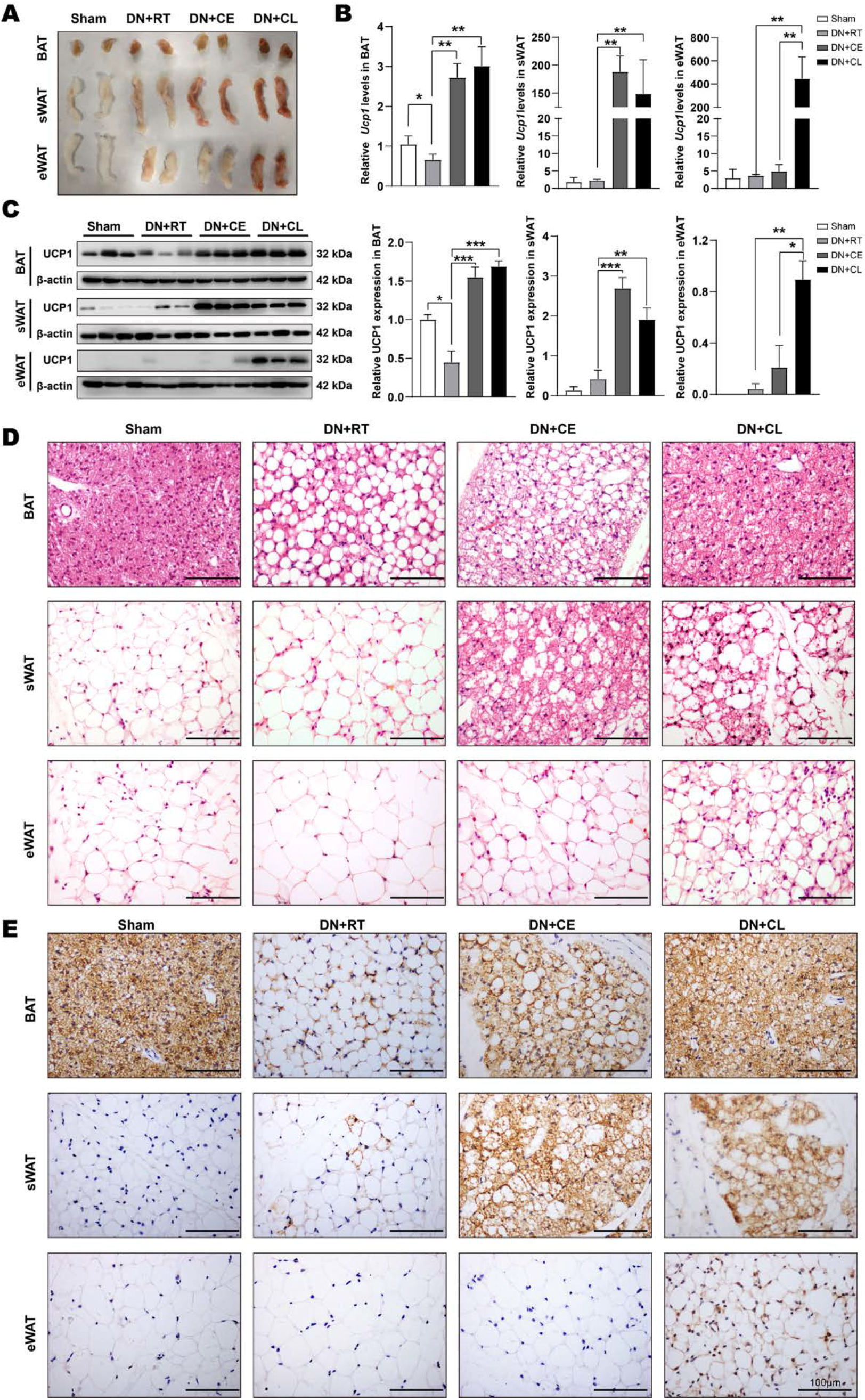
The characteristics of BAT and beige adipocytes in denervated mice after cold exposure or CL316243 injections. (A) The images showing the morphology of BAT (upper), sWAT (middle) and eWAT (lower) in Sham, DN+RT, DN+CE and DN+CL groups. (B) Quantitative realtime PCR showing relative mRNA levels of UCP1 and (C) western blot showing relative UCP1 protein levels in BAT, sWAT and eWAT in Sham, DN+RT, DN+CE and DN+CL groups. (C) Representative sections showing morphological changes and (D) representative immunostaining sections showing UCP1 positive multiocular adipocytes in BAT (upper), sWAT (middle) and eWAT (lower) in Sham, DN+RT, DN+CE and DN+CL groups. Scale bar =100 μm for C and D. A one-factor ANOVA followed by a Tukey’s test was used to make group comparisons. *p < 0.05, **p < 0.01, ***p < 0.001 versus indicated groups. BAT, brown adipose tissue; CE, cold exposure; CL, CL316243; DN, denervation; eWAT, epididymal white adipose tissue; RT, room temperature; UCP1, uncoupling protein 1; sWAT, subcutaneous white adipose tissue.

In denervated BAT, lots of adipocytes were filled with large lipid droplets (whitening), whereas the majority of adipocytes contained smaller lipid droplets after cold exposure and almost all adipocytes showed multilocular lipid droplets morphology after CL316243 injections (Figure 2D). In addition, a number of multiocular adipocytes were present in sWAT of denervated mice upon cold exposure and CL316243 intraperitoneal administration. Moreover, the multiocular adipocytes in eWAT were only observed in denervated mice after CL316243 injections (Figure 2D). No obvious morphological changes were observed in sWAT and eWAT of mice after surgical denervation of BAT. Immunostaining results showed that robust UCP1 positive cells were distributed throughout the entire BAT in Sham, DN+CE and DN+CL groups (Figure 2E). In addition, lots of UCP1-positive multiocular adipocytes (so-called beige adipocytes) were observed and distributed throughout a large area of sWAT in DN+CE and DN+CL groups (Figure 2E). In contrast a few UCP1-postitive cells were present in sWAT in DN+RT group. A number of UCP1-positive beige adipocytes in eWAT were only found in DN+CL, but not in DN+CE group (Figure 2E). These results confirm the beige adipocytes depot in sWAT and eWAT of denervated mice after cold exposure and CL316243 injections.

As showed in Figure 1D, cold exposure increased the levels of food intake in denervated mice, but not body weight at 1 week and 2 weeks. Whereas, CL316243 treatments had no influence on food intake and body weight in denervated mice. The blood glucose levels were significantly improved in DN+CE (Figure 1F, 8.06 ± 0.25 mmol/L) and DN+CL groups (Figure 1F, 6.29 ± 0.36 mmol/L) when compared with the DN+RT group (Figure 1F, 9.34 ± 0.46 mmol/L). Moreover, A one-factor ANOVA showed that the AUC of IPGTT and IPITT were significantly decreased in DN+CE and DN+CL groups when compared with DN+RT group (Figure 1F and G). These data suggest that beige adipocytes in sWAT and eWAT can improve glucose intolerance in denervated mice.

### Beige adipocytes were correlated with up-regulation of Glut1 and Glut4 gene expression in sWAT and eWAT of denervated mice

Glucose transporters are vital membrane-embedded proteins for the uptake of glucose into the cells. Therefore we next examined whether beige adipocytes improved glucose metabolism via upregulation of glucose transporters gene expression. Levels of Glut1 and Glut4 mRNA were significantly decreased in sWAT, eWAT and BAT of mice in DN+RT group when compared with Sham group (Figure 3A). Significant levels of Glut1 and Glut4 mRNA were increased by 2-3 fold in sWAT, eWAT and BAT of mice in DN+CL group when compared with DN+RT group (Figure 3A). There was no significant difference in Glut1 and Glut4 mRNA levels in sWAT, eWAT and BAT of mice between DN+CE and DN+RT group (Figure 3A). Western blot analysis showed that the significant levels of Glut1 and Glut4 proteins were present in sWAT, eWAT of mice in DN+CL group relative to those in DN+RT group (Figure 3B and C). Glut1 and Glut4 proteins levels were significantly decreased in the sWAT, eWAT of mice in the DN+RT group relative to those in Sham group. There was no significant difference in Glut1 and Glut4 proteins levels in sWAT, eWAT of mice between DN+CE and DN+RT group (Figure 3B and C). In the denervated BAT, a significantly decreased levels of Glut1 and Glut4 proteins were detected in DN+RT group (Figure 3D). More importantly, Glut1 and Glut4 proteins levels were also significantly increased in DN+CL group when compared with DN+RT group (Figure 3D). These results indicate that beige adipocytes can improve the glucose intolerance via up-regulation of Glut1 and Glut4 expression in denervated mice.

**Figure 3.**
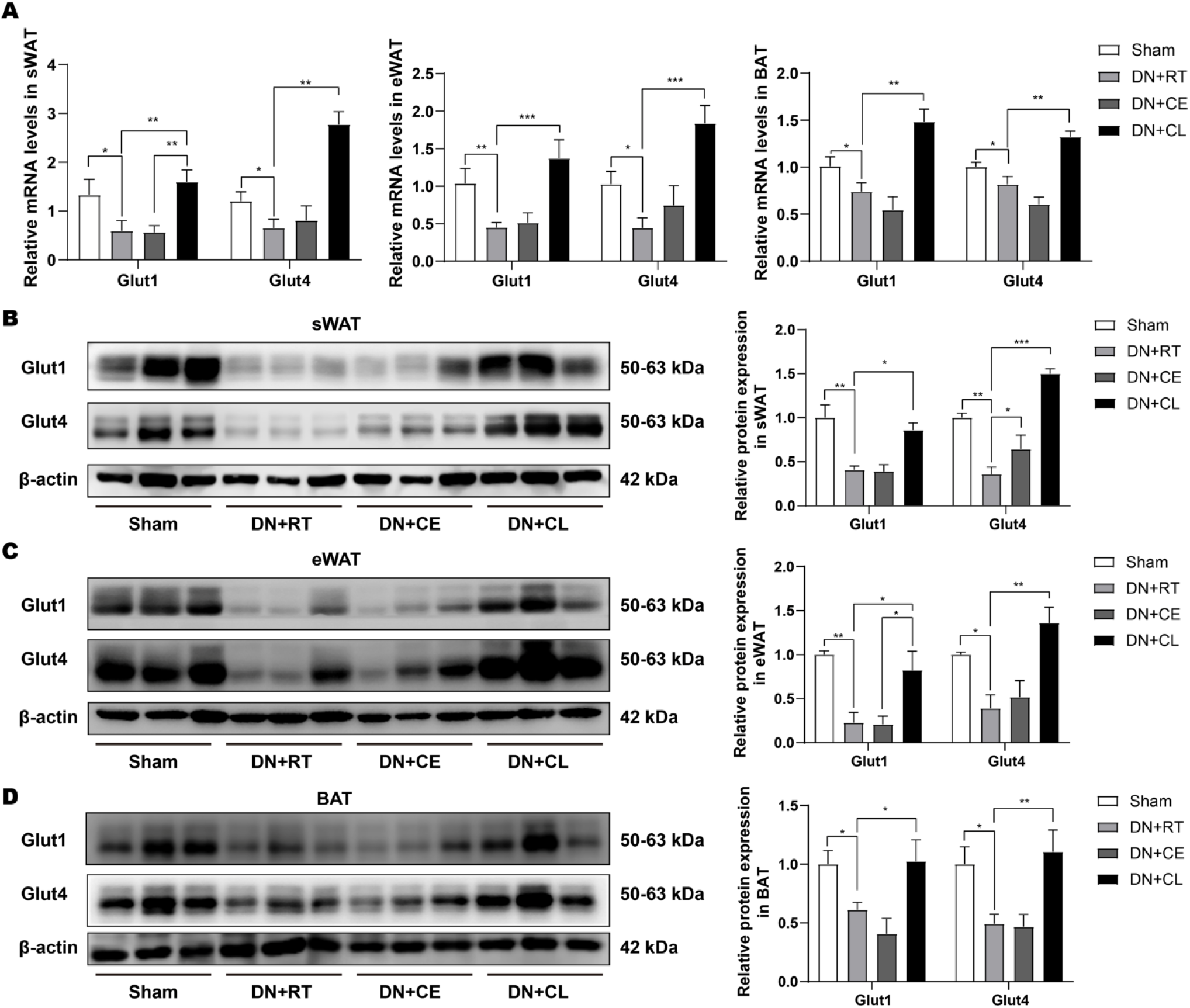
Glucose transporter genes expression in BAT, sWAT and eWAT in denervated mice after cold exposure or CL316243 injections. (A) Quantitative Real-time PCR showing relative mRNA levels of Glut1 and Glut4 in BAT, sWAT and eWAT in denervated mice after cold exposure or CL316243 injections. (B) Western blot showing the protein levels of Glut1 and Glut4 in BAT, sWAT and eWAT in denervated mice after cold exposure or CL316243 injections. A one-factor ANOVA followed by a Tukey’s test was used to make group comparisons. *p < 0.05, **p < 0.01, ***p < 0.001 versus indicated groups. BAT, brown adipose tissue; CE, cold exposure; CL, CL316243; DN, denervation; eWAT, epididymal white adipose tissue; Glut1, glucose transporter 1; Glut4, glucose transporter 4; RT, room temperature; sWAT, subcutaneous white adipose tissue.

### Beige adipocytes independently improved glucose metabolism of mice after removal of BAT

In denervated mice upon cold exposure or CL316243 injections, we observed that denervated BAT showed increased levels of UCP1, Glut1 and Glut4 expression (Figure 2C, 3A and 3D). We can not exclude the contribution of denervated BAT to the improvement of impaired glucose metabolism. Therefore we next did surgical removal of bilateral BAT to examine the specific functions of beige adipocytes independently (Figure 4A). A significant decrease was detected in the food intake of mice after removal of BAT. Whereas, cold exposure or CL316243 treatment promoted the food intake of mice after removal of BAT (Figure 4B). No difference was observed in the body weight of mice in other groups at 0 week, 1 week and 2 weeks (Figure 4C). There was a significant increase of the blood glucose levels in the in RM+RT group (Figure 4D, 9.34 ± 1.30 mmol/L) when compared with Sham group (Figure 4D, 6.43 ± 1.15 mmol/L). The blood glucose levels were significantly decreased in DN+CE (Figure 4D, 6.29 ± 1.14mmol/L) and DN+CL groups (Figure 4D, 8.06 ± 0.83 mmol/L) when compared with that in DN+RT group (Figure 4D, 9.34 ± 0.46 mmol/L). Both AUC of IPGTT and IPITT were significantly decreased in DN+CE and DN+CL groups when compared with those in DN+RT group (Figure 4D and E).

**Figure 4.**
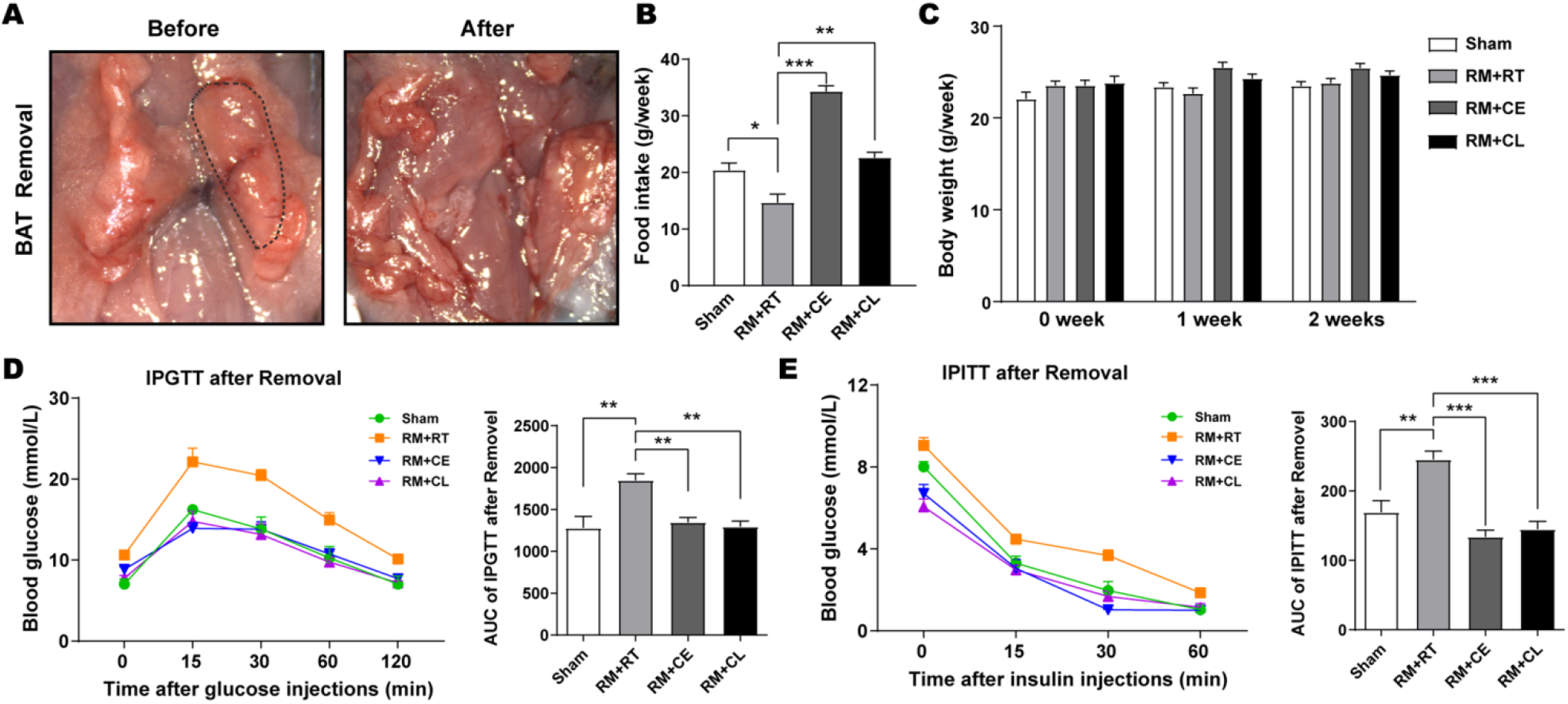
Beige adipocytes improved impaired glucose tolerance of BAT-removal mice. (A) Surgical removal of BAT (dotted circles showing the intact BAT. (B and C) The diagram showing the food intake (B) and body weight at 0 week, 1 week and 2 weeks (C) of mice in Sham, RM+RT, RM+CE and RM+CL groups. (D and E) The diagrams showing the blood glucose (mmol/L) at indicated time points after glucose injections (D, left), AUC analysis during the measurement (D, right), the blood glucose (mmol/L) at indicated time points after insulin injections (E, left) and AUC analysis during the measurement (E, right) of mice in Sham, RM+RT, RM+CE and RM+CL groups. A one-factor ANOVA followed by a Tukey’s test was used to make group comparisons. *p < 0.05, **p < 0.01, ***p < 0.001 versus indicated groups. BAT, brown adipose tissue; CE, cold exposure; CL, CL316243. AUC, area under curve; CE, cold exposure; IPGTT, intraperitoneal glucose tolerance test; IPITT, intraperitoneal glucose tolerance test; RM. Removal; RT, room temperature.

We found that BAT removal did not lead to morphological changes in sWAT and eWAT (Figure 5A). The multiocular adipocytes and robust UCP1 positive cells were distributed throughout the tissue of sWAT and eWAT in mice of RM+CE and RM+CL groups (Figure 5A and B). And no UCP1 proteins were detected in sWAT and eWAT in mice after BAT removal (Figure 5C). UCP1 protein and mRNA levels were significantly increased in sWAT and eWAT in mice of RM+CE and RM+CL groups when compared with those in RM+RT and Sham groups (Figure 5C and D). The browning-related genes including *Pgc1α, Prdm16, Tmem26, CD137, Ppar-α* and *Cidea* were also significantly increased in sWAT in mice of RM+CE and RM+CL groups when compared with RM+RT and Sham groups (Figure 5E). The levels of *Tmem26, CD137, Ppar-α* and *Cidea* mRNA expression were significantly enhanced in eWAT in mice of RM+CL group when compared with RM+RT group (Figure 5F). The levels of *Prdm16 and Cidea* mRNA expression were significantly enhanced in eWAT in mice of RM+CE group when compared with RM+RT group (Figure 5F).

**Figure 5.**
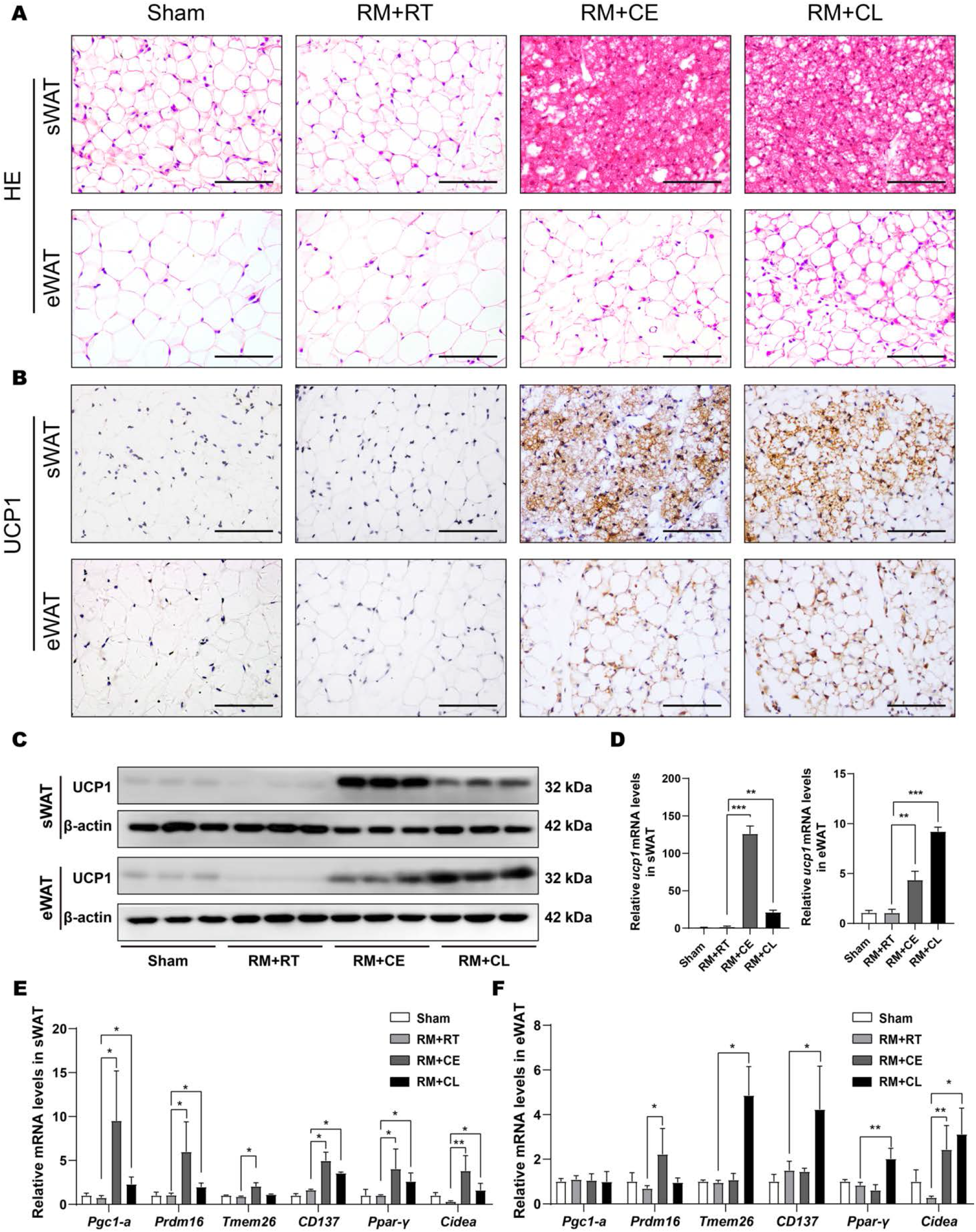
The characteristics and browning related genes expression of Beige adipocytes in BAT-removal mice after cold exposure or CL316243 injections. (A) Representative sections showing morphological changes and (B) representative immunostaining sections showing UCP1 positive multiocular adipocytes in sWAT and eWAT in Sham, RM+RT, RM+CE and RM+CL groups. Scale bar =100 μm for A and B. (C) Western blot showing relative UCP1 protein levels and (D) Quantitative real-time PCR showing relative UCP1 mRNA levels in sWAT and eWAT in Sham, RM+RT, RM+CE and RM+CL groups. (E and F) The diagrams showing browning related genes expression *(Pgc1α, Prdm16, Tmem26, CD137, Ppar-γ* and *Cidea)* in sWAT (E) and eWAT (F) in Sham, RM+RT, RM+CE and RM+CL groups. *p < 0.05, **p < 0.01, ***p < 0.001 versus indicated groups. CE, cold exposure; CL, CL316243; eWAT, epididymal white adipose tissue; RM. removal; RT, room temperature. UCP1, uncoupling protein 1; sWAT, subcutaneous white adipose tissue.

### Beige adipocytes altered energy expenditure of mice after removal of BAT

To determine whether beige adipocytes altered energy expenditure of mice after BAT removal, we further placed the mice after BAT removal injected with CL316243 or saline in metabolic cages to assess the rates of energy expenditure. Although a slight increase in O_2_ consumption (VO_2_) (Figure 6A), CO_2_ production (VCO_2_) (Figure 6B) and energy expenditure (Figure 6C) was noted in RM+RT group relative to those in Sham group, no significant difference was observed between these two groups. O_2_ consumption (Figure 6A), CO_2_ production (Figure 6B) and energy expenditure (Figure 6C) in BAT-removal mice were enhanced by CL316243 injections. In addition, the data for respiratory quotient presented a significant decrease in RM+RT and RM+CL groups when compared to that in Sham group (Figure 6D). Pedometer data showed a significant enhancement at daytime, nighttime, and 24 hours in RM+RT group when compared to that in RM+CL and Sham groups (Figure 6E). Similar tendencies were observed in another activity measurement, including the sum of X- and Y-break (Figure 6F). There was no difference in food intake and water drink among these groups (Figure 6G).

**Figure 6.**
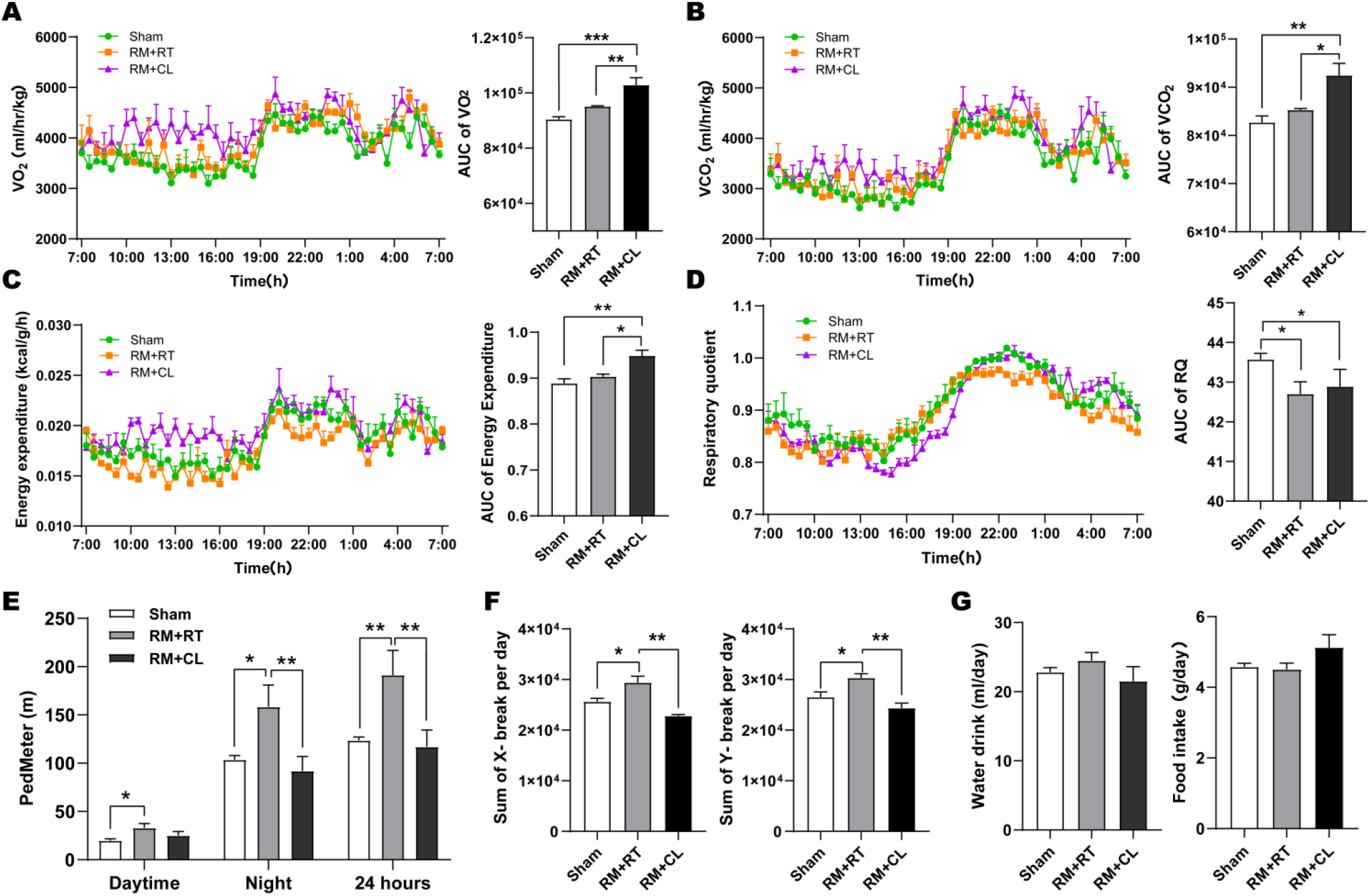
Beige adipocytes altered energy expenditure of mice after removal of BAT. (A) O_2_ consumption of mice in Sham, RM+RT and RM+CL groups (Left); AUC analysis of VO_2_ during the measurement (Right). (B) CO_2_ consumption of mice in Sham, RM+RT and RM+CL groups (Left); AUC analysis of VCO_2_ during the measurement (Right). (C) Energy expenditure of mice in Sham, RM+RT and RM+CL groups (Left); AUC analysis of energy expenditure during the measurement (Right). (D) Respiratory quotient of mice in Sham, RM+RT and RM+CL groups (Left); AUC analysis of energy expenditure during the measurement (Right). (E-G) The diagrams showing pedometer (E), sum of X break (F, left), sum of Y break (F, right), water drink (G, left) and food intake (G, right) of mice in Sham, RM+RT and RM+CL groups. *p < 0.05, **p < 0.01, ***p < 0.001 versus indicated groups. AUC, area under curve; CL, CL316243; RM. removal; RQ, respiratory quotient.

### Up-regulation of glucose transporter genes and activation of insulin signaling in sWAT and eWAT of BAT-removal mice after cold exposure or CL316243 injections

We examined glucose transporter genes expression in sWAT and eWAT of BAT-removal mice after cold exposure or CL316243 injections. There was a significant increase in the levels of Glut1 and Glut4 proteins and mRNA in sWAT and eWAT in RM+CL group when compared with those in RM+RT group (Figure 7A, B and Supplementary S1). No difference was observed in the levels of Glut1 and Glut4 proteins and mRNA in sWAT and eWAT in RM+CE group when compared with RM+RT group (Figure 7A, B and Supplementary S1).

**Figure 7.**
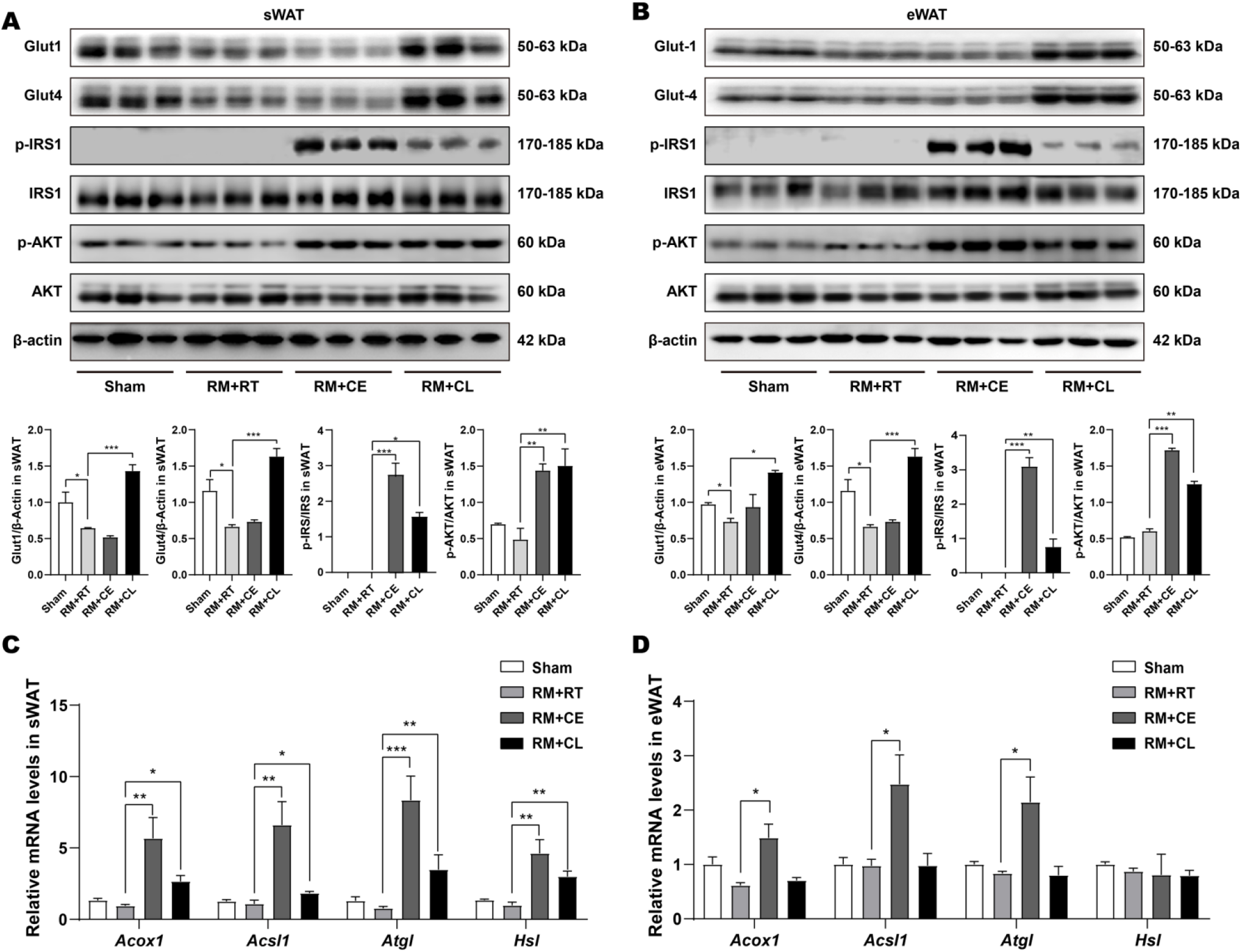
Glucose transporter genes expression and insulin signaling in sWAT and eWAT in BAT-removal mice after cold exposure or CL316243 injections. (A and B) Western blot showing relative Glut1, Glut4, p-IRS1/IRS1, p-AKT/AKT protein levels in sWAT (A) and eWAT (B) in Sham, RM+RT, RM+CE and RM+CL groups. (C and D) The diagrams showing relative lipolytic genes expression *(Acox1, Acsl1, Atgl,* and *Hsl)* in sWAT (C) and eWAT (D) in Sham, RM+RT, RM+CE and RM+CL groups. *p < 0.05, **p < 0.01, ***p < 0.001 versus indicated groups. ACOX1, peroxisomal acyl-coenzyme A oxidase 1; ACSL1, long-chain fatty acid-CoA ligase 1; ATGL, lipolytic enzymes adipose triglyceride lipase; CE, cold exposrue; CL, CL316243; HSL, hormone-sensitive lipase; RM. Removal; RT, room temperature.

Next, we examined insulin signaling in sWAT and eWAT of BAT-removal mice after cold exposure or CL316243 injections. There was no difference in insulin receptor substrate 1 (IRS1) protein in all groups. In contrast, phosphorylation IRS1 (p-IRS1) was significantly increased in sWAT and eWAT of BAT-removal mice after cold exposure or CL316243 injections (Figure 7C and D). Furthermore, the downstream p-AKT was also significantly increased in sWAT and eWAT of BAT-removal mice after cold exposure or CL316243 injections, and no difference was detected in AKT protein among groups (Figure 7A and B).

### Lipolytic related genes expression in sWAT and eWAT of BAT-removal mice upon cold exposure or CL316243 injections

As showed in Figure 7C, the lipolytic related genes expression, including *Acox1, Acsl1, Atgl,* and *Hsl* were significantly increased in sWAT of BAT-removal mice upon cold exposure or CL316243 injections. Whereas, there was no significant difference in lipolytic related genes expression in sWAT and eWAT in RM+RT group when compared with those in Sham group (Figure 7C). *Acox1, Acsl1 and Atgl* genes expression were significantly increased in eWAT in RM+CE group when compared with that in RM+RT. There was no significant elevated expression of lipolytic genes, including *Acox1, Acsl1, Atgl,* and *Hsl* in eWAT in RM+CL group when compared with those in RM+RT group (Figure 7D). These data suggested an increased level of lipolytic activity in white adipose tissue of BAT-removal mice upon cold exposure or CL316243 injections.

## DISCUSSION

The present study firstly shows that beige adipocytes upon challenges have a beneficial effect on glucose metabolism and energy expenditure in the absence of BAT in male mice. The data show that surgical denervation of BAT leads to glucose and insulin intolerance in vivo. Beige adipocytes partly contribute to impaired glucose metabolism because denervated BAT can be activated by cold exposure or CL316243 injections. Our results further reveal that beige adipocytes can independently improve glucose intolerance in the absence of BAT. The insulin signaling was activated in BAT-removal mice upon cold exposure. Both the activation of insulin signaling and upregulation of glucose transporter genes expression were observed in BAT-removal mice with CL316243 treatments. These data show that beige adipocytes induced by cold exposure or β3-adrenoceptor agonist may have different mechanisms to improve glucose and insulin intolerance in vivo. Beige adipocytes can also enhance energy expenditure and lead to a greater lipolysis in sWAT and eWAT in the absence of BAT in vivo.

It has been demonstrated that BAT contributes to glucose metabolism in rodents (20, 22, 23). We therefore hypothesize that BAT ablation may lead to glucose intolerance in mice. Currently, we found the impaired glucose metabolism both in BAT-denervated and BAT-removal animal models, as evidenced by the increased levels of blood glucose and glucose area under curve. A recent study reported that BAT removal did not impair systemic glucose metabolism in lean mice (53), which is contradictory to the current results. Two factors may contribute to this inconsistency. Firstly, this discrepancy may be due to the different housing conditions. The housing temperature of 25°C used by Grunewald’s group limits thermal stress and housing at temperature around 20°C in the present study is able to result in increased adrenergic activity to brown adipose tissue to maintain thermal homeostasis (54). Secondly, the different study duration (3 weeks versus 12 weeks) may also contribute to this contradiction. When BAT is lacking, the compensatory capacity of other fat depots may be a time course-dependent manner. It will be interesting to determine the mechanism to compensate for the loss of functional BAT.

Beige adipocytes are emerging as an interesting issue in obesity and metabolism research (3, 6, 21, 26, 29). More importantly, several groups have reported that brown adipose tissue in human adults appears to be primarily composed of beige adipocytes (15, 16, 24, 25). An important question is whether beige adipocytes have the same functions as brown adipocytes. Based on amounts of UCP1 protein, UCP1 mRNA and oxygen consumption, it would tend to estimate that the maximum capacity of the most recruited beige adipocytes could amount to 25% of the total UCP1-dependent thermogenic capacity (17). In our animal models of BAT ablation generated by surgical removal, we found that beige adipocytes upon cold challenge or CL316243 activation could improve glucose intolerance in the absence of BAT in male mice. We also found that the activity of insulin signaling related to glucose transporter translocation was markedly increased in white adipose tissue of BAT-removed mice under cold exposure or CL316243 treatments. In addition, glucose transporter genes expression (Glut1 and Glut4) were also greatly increased in white adipose tissue of BAT-removed mice upon CL316243 treatments. These results suggest that beige adipocytes could contribute to impaired glucose metabolism when BAT is lacking. Seale’s group has reported a favorable rise in whole-body energy expenditure in the aP2-PRDM16 transgenic mouse with a selective browning of sWAT, which suggest that beige adipocytes may contribute to the overall metabolic phenotype observed in this transgenic mouse (55). Similarly we also observed increased metabolic efficiency in BAT-removal mice with CL316243 treatments, as evidenced by the enhancement of oxygen consumption, carbon dioxide production and motor activity. These findings indicate that beige adipocytes could contribute to glucose metabolism and energy expenditure in the absence of BAT. A recent study reported that bilateral denervation of BAT could not increase iWAT oxidative metabolic activity (48), which is inconsistence with our results. This discrepancy may be due to housing temperature (4°C versus. 25°C) and the animal model (denervation versus removal). Nevertheless, the specific functions of beige adipocytes still need to be further explored.

Several lines of evidence have demonstrated the difference of beige adipocytes formation in sWAT and eWAT in response to cold exposure and β3-adrenoceptor agonist stimulation (27, 32, 33, 35, 56, 57). Wang and colleagues have reported that beige adipocytes generated by CL316243 and rosiglitazone injections have different genetic signatures (58). Therefore, beige adipocytes generated by different stimuli may have distinct mechanisms to exert functions. We observed the improvement of glucose intolerance in BAT-removal mice upon cold or CL316243 treatments simultaneously. We found that both protein levels of p-IRS1, p-AKT and glucose transporter genes were greatly increased in sWAT and eWAT upon CL316243 treatments. Whereas, protein levels of p-IRS1 and p-AKT, but not glucose transporter genes, were significantly enhanced in sWAT and eWAT of mice upon cold exposure. These results show that beige adipocytes induced by cold exposure or β3-adrenoceptor agonist stimulation may have different mechanisms to improve glucose intolerance in vivo. Under cold conditions, in addition to the contribution of beige adipocytes, the glucose uptake could be also mediated by skeletal muscles metabolic activation even when shivering is minimized (39, 59). It will be of interest to determine the different functions of beige adipocytes under different stimuli. And it also needs to identify the beneficial effects of beige adipocytes in WAT on human obesity and associated metabolic disorders.

In rodents, interscapular BAT has sympathetic and sensory system innervation (60). Sympathetic innervation plays a critical role in energy metabolism and thermogenesis of BAT via UCP1 activity. It has been reported that chemical and surgical denervation are selected to produce sympathectomy. Chemical Injections of guanethidine (blocking sympathetic conduction) or 6-OHDA (a catecholaminergic neurotoxin) are restrictive due to the incomplete elimination of sympathetic nerves (50). Currently, an ablation of nerve bundles of bilateral BAT was used to abolish the function of BAT in vivo. In consistence with previous studies (48, 49, 61), the surgical denervation was confirmed by the significant elimination of TH in BAT in the current study. We found that whitening occurred in denervated of BAT as evidenced by the uniocular morphology and decreased levels of UCP1 protein. Whereas, the majority of uniocular adipocytes in denervated BAT regained their multiocular morphology and robust UCP1 immunostaining under chronic cold conditions (4°C) and β3-adrenoceptor agonist treatments (CL316243). In addition, UCP1 gene expression in denervated BAT was enhanced upon cold exposure and CL316243 injections. Glucose transporter genes were also increased in denervated BAT of mice after CL316243 treatments. These results are the first to suggest that the function of BAT is temporarily abrogated by surgical denervation, and the denervated BAT is prone to be replenished by external stimuli.

BAT has the capacity to dissipate energy via the activity of UCP1 and WAT is able to store energy in the form of triglycerides. The subcutaneous ingWAT (denoted as sWAT or iWAT) is commonly regarded as the white adipose depot with the strongest browning capacity (62). An interesting issue is what occurs in sWAT when BAT is lacking. A recent study has reported that bilateral denervation of BAT can promote sWAT browning at thermoneutrality (48). However, A more recent study showed that no browning of sWAT was detectable in BAT-denervated mice (61). Similarly, in BAT-denervated and BAT-removal mice at room temperature, we did not achieve significant evidence for browning of sWAT. No apparent morphological alterations of adipocytes, no widespread UCP1 staining, and no significant increase in UCP1 protein and mRNA levels were detected in sWAT of BAT-denervated and BAT-removal mice at room temperature in our study. We did not achieve evidence for enhancement of lipolytic activity in sWAT in BAT-removal mice at room temperature. Schulz and colleagues have claimed that denervation of BAT triggered a greater browning of WAT under conditions of CL316243 injections (51). Currently, our study provides several lines of evidence that surgical removal of BAT results in an extensive browning of sWAT in response to cold exposure and CL316243 treatments. These results are in good consistent with a study showing beige adipocytes formation was enhanced upon BAT removal and cold exposure (63).

In summary, our data show the beneficial effect of beige adipocytes upon cold challenge or CL316243 activation in the absence of BAT. The data show that beige adipocytes induced by external stimuli can partly improve impaired glucose metabolism in denervated mice. However, we firstly show that the function of the denervated BAT is temporarily inhibited and could be activated by cold exposure or β-adrenoceptor agonist administration. We further confirm that surgical removal of BAT results in glucose intolerance in male mice. Beige adipocytes upon cold challenge or CL316243 activation can independently improve glucose intolerance when BAT is lacking. Activation of insulin signaling and increased levels of glucose transporter genes may constitute the different mechanisms that beige adipocytes improve impaired glucose metabolism. Our study supports the treatments selectively targeting white adipose tissue to generate beige adipocytes of clinical trials on obesity and associated metabolic disorders.

## ACKNOWLEDGMENTS

We thank Xi-wen Xiong professor for help with energy metabolic analysis.

## GRANTS

This study was supported by National Natural Science Foundation of China (81602532), Natural Science Foundation of Beijing Municipality(5202004), Beijing Municipal Organization Department Talents Project (2015000020124G113), Support Project of High-level Teachers in Beijing Municipal Universities in the Period of 13th Five-year Plan (IDHT20170516).

## DISCLOSURES

The authors declare no conflicts of interest, financial or otherwise.

